# Lactate Dehydrogenase Activity and Carbohydrate Metabolism under Vanadium Citrate Exposure: Sex- and Dose-Dependent Effects in Rat Tissues

**DOI:** 10.64898/2026.08.23.746541

**Authors:** Ruslana Iskra, Halyna Klymets, Iryna Oliynyk

## Abstract

Vanadium (V) is a potential insulinomimetic that can modulate carbohydrate metabolism, but its biological effects are sensitive to chemical form, concentration, and sex. Chelation of vanadium with organic ligands, in particular citrate, allows to increase its bioavailability and optimize pharmacokinetic properties. The aim of the study was to evaluate tissue-, dose-, and sex-dependent changes in physiological parameters and activity of the key glycolytic enzyme — lactate dehydrogenase (LDH) — under the influence of vanadium citrate. The study was conducted on 6-week-old Wistar rats of both sexes. The animals received vanadium citrate orally for 36–38 days at doses of 3, 12.5, and 50 μg VCit/kg body weight. LDH activity in skeletal muscle, liver, kidney, and pancreas was investigated. No pronounced toxic effect on physiological parameters was detected: body weight dynamics corresponded to age norms, no behavioral changes were observed. LDH activity demonstrated pronounced sexual dimorphism and depended on the dose received. It was established that the optimal dose, which provides a modulating effect without signs of metabolic stress, for females is 12.5 μg VCit/kg, while for males - 3 μg VCit/kg. The most significant changes in LDH activity were recorded in the pancreas at a dose of 50 μg V/kg, where the indicators decreased from 0.81 to 0.31 μmol/(min×mg protein) in females and from 1.02 to 0.28 μmol/(min×mg protein) in males.

The effect of vanadium citrate on carbohydrate metabolism, as well as its dose-, tissue-and sex-specific nature, is likely determined by a dual action: the insulin-like effect of vanadium (redirecting pyruvate to oxidation) and the allosteric inhibition of glycolysis by the citrate ligand (substrate limitation for LDH). The obtained results emphasize the importance of considering sex and dose in the research and development of metabolically active compounds.

## Introduction

Vanadium (V) has attracted considerable attention as a biologically active trace element capable of modulating carbohydrate metabolism due to its insulin-like properties [1, 2]. It is known that its compounds can mimic phosphate groups, affecting the activity of protein tyrosine phosphatases and regulating key signaling cascades associated with glucose metabolism. As a result, vanadium is able to change cellular glucose uptake, glycolytic flux and the direction of pyruvate utilization. The biological effect of vanadium largely depends on its chemical form. Chelation with organic ligands, in particular citrate, increases its bioavailability and changes its pharmacokinetic properties. Citrate was selected as the chelating ligand because, unlike purely transport-oriented ligands (e.g., sulfate or simple organic acids), it is itself a bioactive tricarboxylic acid cycle intermediate with allosteric regulatory activity toward phosphofructokinase-1 [3]. This dual functionality — as both a bioavailability-enhancing carrier and an independent metabolic modulator — distinguishes vanadium citrate from other vanadium compounds (e.g., vanadyl sulfate, sodium metavanadate) previously used in metabolic studies, and provides the rationale for its selection in this work. At the same time, citrate is not only a transport component, but also an active metabolic regulator. As an intermediate of the tricarboxylic acid cycle, it acts as an allosteric inhibitor of phosphofructokinase-1, a key enzyme in glycolysis, allowing it to limit glycolytic flux and alter the availability of substrates for subsequent metabolic transformations [3] (see Fig 1).

**Fig 1.**
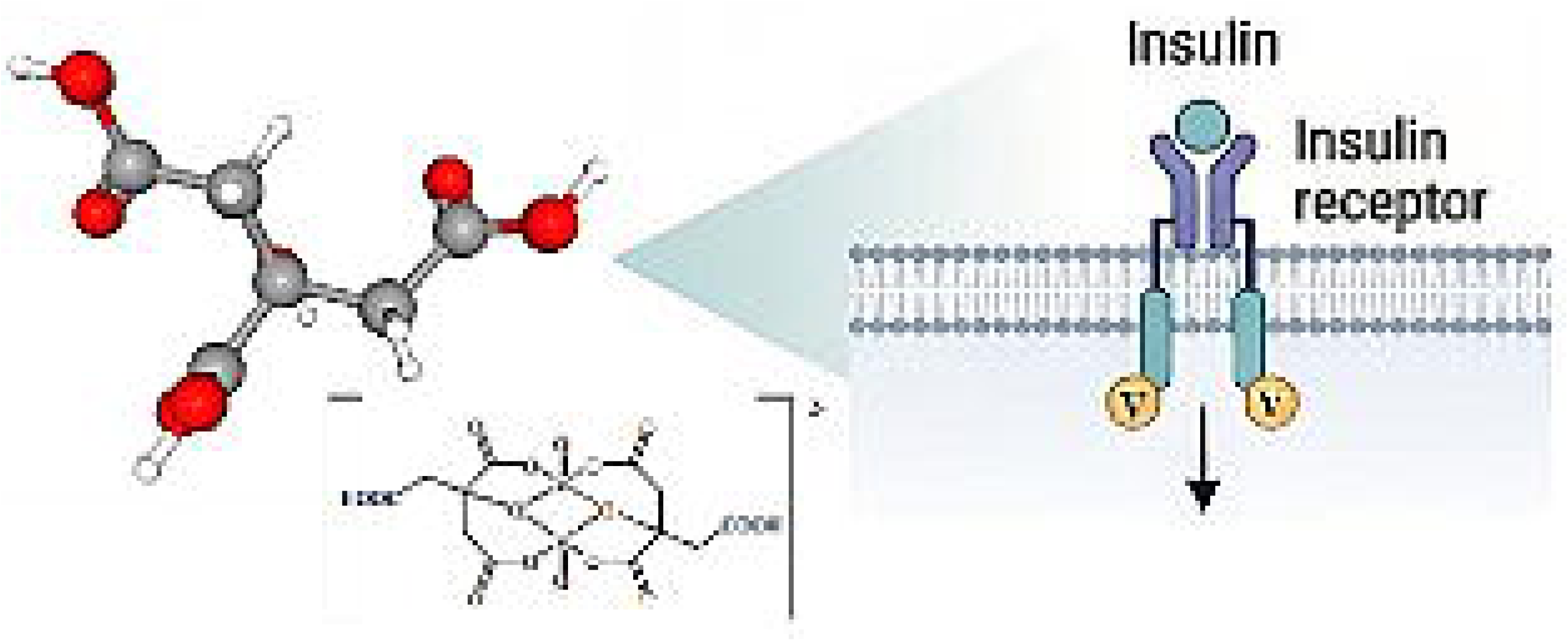
Molecular structure of the vanadium citrate complex (after Tsaramyrsi M. [3]) and its proposed insulin-like mode of action based on modulation of insulin receptor signalling pathways.

Thus, the vanadium–citrate complex potentially implements a dual mechanism of action [4]: on the one hand, through the insulinomimetic effect of vanadium, which promotes glucose utilization, and on the other hand, through the regulatory effect of citrate on glycolysis [5]. However, the integrated effect of such interaction at the level of tissue metabolism remains poorly understood. Lactate dehydrogenase (LDH; EC 1.1.1.27) is a key enzyme that catalyzes the interconversion of pyruvate and lactate and plays a central role in maintaining cellular redox balance. The enzyme exists in the form of five isoforms (LDH –LDH), which are formed by different combinations of the LDH-A (M) and LDH-B (H) subunits and are characterized by pronounced tissue specificity.

LDH-B subunit-rich isoforms (LDH□–LDH□) predominate in aerobic tissues such as the heart and kidney, where they promote the oxidation of lactate to pyruvate for subsequent incorporation into mitochondrial oxidation. In contrast, LDH-A-containing isoforms (LDH□–LDH□) dominate in glycolytic tissues, particularly skeletal muscle and liver, where they promote the reduction of pyruvate to lactate and maintain glycolytic flux.

This functional specialization of LDH isoforms determines the tissue-specific direction of pyruvate metabolism - either towards anaerobic glycolysis or towards aerobic oxidation. Accordingly, changes in LDH activity may reflect not only the overall intensity of glycolysis, but also a shift in the metabolic balance between lactate generation and mitochondrial oxidation.

Considering that vanadium exhibits insulinomimetic properties and is able to influence glucose utilization, and citrate acts as an allosteric inhibitor of phosphofructokinase-1, it can be assumed that the vanadium–citrate complex is able to modulate LDH activity both by changing the glycolytic flux and by redirecting pyruvate to oxidative pathways. However, the nature of these changes may differ significantly depending on the tissue isoform structure of the enzyme.

Despite available data on the biological activity of vanadium compounds, their tissue-, dose-, and sex-specific effects on LDH activity under normal physiological conditions remain insufficiently characterized. Most prior work has been performed on pathological (e.g., diabetic) models, which confounds primary metabolic effects of vanadium with disease-related metabolic dysregulation, and rarely addresses sex as a biological variable. Given the growing trend of off-label use of metabolic modulators, such as GLP-1 receptor agonists, for weight management in otherwise healthy individuals [6], characterizing the physiological effects of insulinomimetics such as vanadium citrate in an intact organism is of particular relevance.

### Objectives

The aim of this study was therefore to evaluate tissue-, dose-, and sex-dependent changes in LDH activity in healthy Wistar rats under long-term vanadium citrate exposure, in order to (i) identify early, sex-specific signatures of its effect on carbohydrate metabolism, and (ii) assess potential dose-dependent metabolic risks in the absence of underlying pathology.

## Materials and methods

### Animals and conditions of detention

The studies were performed on healthy Wistar male and female white laboratory rats weighing 140-160 g aged 6 weeks (early puberty (sexual differentiation)). The choice of 6-week-old rats was due to the study of metabolism during the period of active puberty and the formation of hormonal levels. The selected age corresponds to the early pubertal period, which was intentionally chosen to capture sex-specific metabolic programming rather than fully stabilized adult hormonal profiles. Male and female were divided into four groups of 6 animals each (total 48 animals). The number of animals per group (n=6) was determined to ensure sufficient statistical power (≥80%) to detect the expected biological effects of vanadium citrate, based on data from previous studies. This sample size also complied with the ethical principles of 3R (Reduction), minimizing the use of animals.

Rats were housed in standard polypropylene cages (2 animals per cage with a partition for the period of nocturnal activity) under controlled temperature (22 ± 2□) and lighting regime (12 h light/12 h dark) in a vivarium and had free access to standardized commercial food and drinking water.

### Compound administration schedule and dosage

Water (Group I) and vanadium citrate solutions (Groups II–IV) were administered orally using a controlled drinking protocol to ensure accurate dosing. Animals in experimental groups II, III, and IV received vanadium citrate solutions corresponding to daily vanadium citrate doses of 3, 12.5, and 50 μgVCit/kg body weight, respectively.

Every evening at 8:00 PM, animals received a fixed volume of 10 mL per 100 g body weight of either drinking water (control group) or vanadium citrate solution (experimental groups) of the appropriate dosage. Considering the body weight of the animals (140–160 g), this volume ensured the full consumption of the calculated dose during the 12 hours of nocturnal activity of the animals.

The following morning at 8:00 AM, the water and solution bottles were checked:

- if the entire volume was consumed, the animals were given water ad libitum;
- if the solution was not completely consumed, the animals were observed until complete consumption, after which they were allowed free access to water.

This approach ensured that each animal received the prescribed daily dose of vanadium citrate, while minimizing variability associated with individual differences in spontaneous water consumption.

The body weight of the animals was measured every 5 days to maintain accurate dosing of vanadium citrate per kilogram of body weight and volume of fluid given overnight throughout the experiment.

Animals were euthanized on day 36-38 by decapitation under thiopental anesthesia, administered in a dose sufficient to induce deep surgical anesthesia [7]. The period was chosen as sufficient for the formation of stable metabolic adaptation to chronic intake of vanadium, which corresponds to long-term (many years - 3–4 years of human life during the period of active youth) exposure in humans. Therefore, the animals did not experience pain during the study. Estrous cycle staging was not performed; however, randomization and group housing were applied to minimize cycle-related bias, and the study design reflects an integrated physiological response rather than phase-specific effects.

### Preparation of tissue homogenates

Ten percent homogenates of skeletal muscle, kidney, liver and pancreas of animals were prepared for further studies. Homogenization was performed in an appropriate buffer (10 ml), using a type 302 homogenizer (Warsaw, Poland) and tissue samples weighing 1 g.

Homogenization of tissue samples was performed in an isotonic Tris-based buffer consisting of 50 mM Tris-HCl (pH 7.4), supplemented with 0.25 M sucrose to maintain osmotic stability. EDTA (1 mM) was included to chelate divalent metal ions and reduce metal-dependent protease activity, while DTT (1 mM) was added to protect thiol groups and preserve enzyme activity. A protease inhibitor cocktail (1×) was added immediately prior to homogenization.

Tissue samples were placed in a pre-cooled vessel and homogenized to obtain a homogeneous mass with the maximum possible degree of grinding.

### Determination of LDH activity

LDH activity was determined spectrophotometrically based on the oxidation of NADH to NAD□ in the presence of pyruvate, according to the method described by Kendig (2007) [8] with minor modifications. Measurements were performed using a UNICO 1205 spectrophotometer (USA) at 37°C by monitoring the decrease in absorbance at 340 nm over a 5-minute period. Enzyme activity was assayed in 0.2 M Tris-HCl buffer (pH 7.5), with a total reaction volume of 3 mL. Tissue homogenates were used immediately for LDH activity measurements. LDH activity was normalized to the total protein concentration determined by the Bradford method in μmol/(min×mg of protein).

For the research, vanadium citrate (Vanadium Citrate) was used, obtained using a nanotechnological synthesis method as described in [9]. This method yields a stable aqueous colloidal solution of vanadium (IV) aquachelate with a stock concentration of 1 g/L, which was freshly diluted to the required doses immediately before administration. Importantly, the preparation does not contain unreacted metallic vanadium or vanadium oxide nanoparticles; therefore, the administered dose corresponds exclusively to biologically active vanadium (IV) aquachelate [10].

Laboratory animals were treated in accordance with the standards of the European Convention for the Protection of Vertebrate Animals, used for research and scientific purposes (Strasbourg, 1986). Protocol of the Bioethics Committee meeting at the Institute of Animal Biology No. 89 dated July 8, 2023.

### Calculation of the stability index

To assess the overall stability of the body’s metabolic reactions to the introduction of vanadium citrate, a stability index (SI) was introduced. This indicator was defined as the average absolute deviation of LDH activity from control values (taken as 100%) in all studied tissues.

The control group, respectively, corresponds to the state of physiological homeostasis. Therefore, any deviation in enzyme activity (both increase and decrease) was considered by us as a sign of “destabilization” or restructuring of the metabolic profile under the influence of the studied compound.

For each dose, SI was calculated using the formula “Metabolic Deviation Index”:

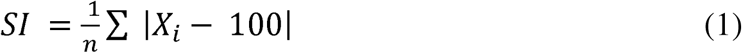

where: *X_I_* — % of control for each tissue, n — number of tissues.

In the appropriate interpretation, a low SI value indicates that tissue metabolism remains close to the control norm, while a high value indicates a pronounced systemic response or potential metabolic stress. Note that SI has certain methodological limitations that should be considered when interpreting: SI depends on which organs are included in the analysis. Since different tissues have different metabolic roles (e.g., liver as a center of gluconeogenesis, versus muscle as a glucose consumer), SI reflects the systemic response, not the specificity of an individual organ. The formula uses the absolute value of the deviation | *X_i_* - 100|. This means that inhibition of the enzyme by 20% and its activation by 20% will make the same contribution to “destabilization”. In a biological sense, these processes may have different meanings: for example, a moderate decrease in LDH in the muscles may indicate a transition to more efficient oxygen oxidation (positive adaptation), while a sharp decrease in the kidneys may be a sign of toxicity.

But at the same time, by calculating SI, it is possible to determine doses with minimal side effects on a healthy organism. However, for models of pathology (for example, diabetes), the “ideal” may not be the minimum SI value, but the one that ensures the return of the changed indicators to normal. Note that for the correct calculation of SI, it is important to have homogeneous groups of animals, since large individual variability within one group may artificially overestimate the index.

### Statistical analysis

Interference statistical analysis included determination of the arithmetic mean value and corresponding confidence intervals of each indicator in the groups. Two parallel measurements were performed for each sample; if the results differed by more than 5%, two additional measurements were performed. Data points affected by technical artifacts or inappropriate repeated measurements were excluded from further analysis.

Statistical processing was carried out using descriptive statistics and analysis of variance methods. Double repetition of measurements allowed calculating the average value of each biochemical indicator of carbohydrate metabolism for each individual sample. All groups were checked for compliance with normal distribution and homogeneity of variances.

Before applying parametric analysis methods (in particular, ANOVA), the data distribution in each group was checked for compliance with normality. For this, the Shapiro–Wilk criterion was used. In the case of multiple comparisons, appropriate Bonferroni corrections were applied.

After checking normality, a test for homogeneity of variances between groups was performed using Levene’s criterion.

Comparison of mean values of physiological parameters between groups was performed using parametric analysis of variance (two-way ANOVA), since the distributions followed a normal law and the variances were homogeneous. Differences were considered statistically significant at p < 0.05.

In addition, to quantify the effect size (Effect Size) and confirm the statistical power of the results, partial eta-square was calculated. The ² estimate allows us to determine what proportion of the total variance of the indicator is explained by the studied factors (gender, dose, and their interaction). Statistical analysis was performed using GraphPad Prism software (version 9.1, GraphPad Software, La Jolla, CA, USA). Data visualization and graphical analysis, including radar charts, were performed using Python (version 3.12.3) with the Matplotlib and Seaborn libraries.

## Results

### General physiological status and body weight dynamics

The doses of vanadium citrate used in the study (3, 12.5 and 50 μgVCit/kg body weight) were well tolerated by the experimental animals during the 36-day experiment. No clinical signs of toxicity, disturbances in food or water consumption, or abnormal behavioral patterns were observed. All groups of animals demonstrated body weight gain, which is the physiological norm for 6-week-old Wistar rats in the phase of active growth and puberty. The absence of pathological weight loss confirms that the selected drug concentrations do not cause pronounced metabolic depletion. Despite the general tendency to increase, the rate of weight gain varied depending on sex and dosage (see Fig 2).

**Fig 2.**
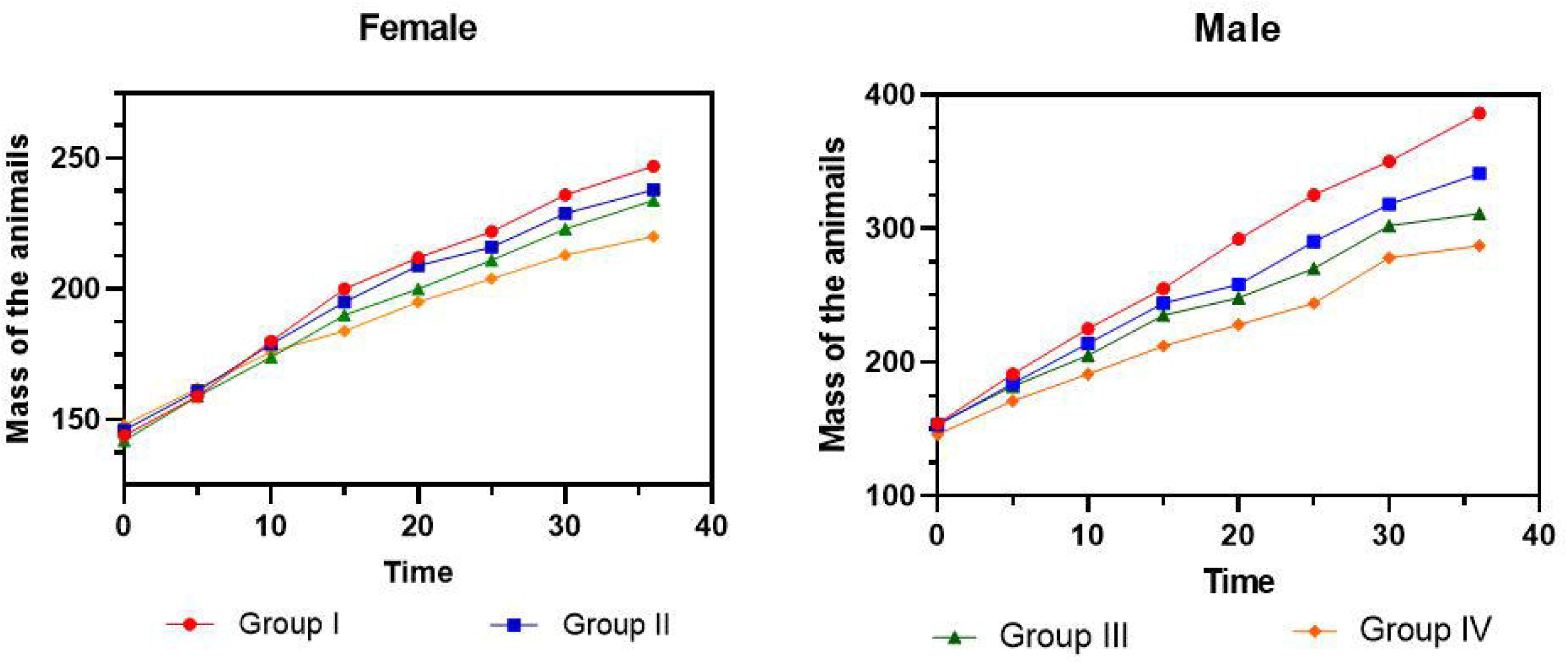
Body weight dynamics of male and female Wistar rats under the influence of different doses of vanadium citrate.

In the female group, on the 35th day of the experiment, body weight gain decreased by 10.68% (at a dose of 3 μgVCit/kg body weight) and by 13.59% (at a dose of 12.5 μgVCit/kg body weight) compared to the control group. In the male group, the inhibitory effect was more pronounced, body weight gain decreased by 19.27% (3 μgVCit/kg body weight) and by 28.4% (12.5 μgVCit/kg body weight) compared to the corresponding control. The administration of vanadium citrate, especially in high doses (50 μgVCit/kg body weight), led to a certain slowdown in weight gain compared to the control group by 30.1% in the female group and by 35.3% in the male group.

### Lactate dehydrogenase activity in skeletal muscle

LDH, an enzyme that plays a crucial role in glycolysis and ATP production in skeletal muscle [11]. In the skeletal muscles of female rats of the control group, LDH activity is significantly lower (8.08 ± 0.5 μmol/(min×mg of protein)) than in males (12.89 ± 0.48 μmol/(min×mg of protein)), which indicates a lower dependence of female muscles on anaerobic glycolysis at rest, compared to males. While in males, glycolysis plays a predominant role in the energy supply of muscles.

Under the influence of vanadium citrate at doses of 3 and 50 μgVCit/kg body weight, a decrease in LDH activity was observed in the skeletal muscles of females by 26.62% and 31.97%, respectively, while the decrease during vanadium consumption at a dose of 12.5 μgVCit/kg body weight was only 7.72% (p = 0.3). Males demonstrated a gradual decrease in enzyme activity under the influence of all citrate doses by 5.81%, 23.03 and 32.55% in groups II, III and IV, respectively (Fig 3).

**Fig 3.**
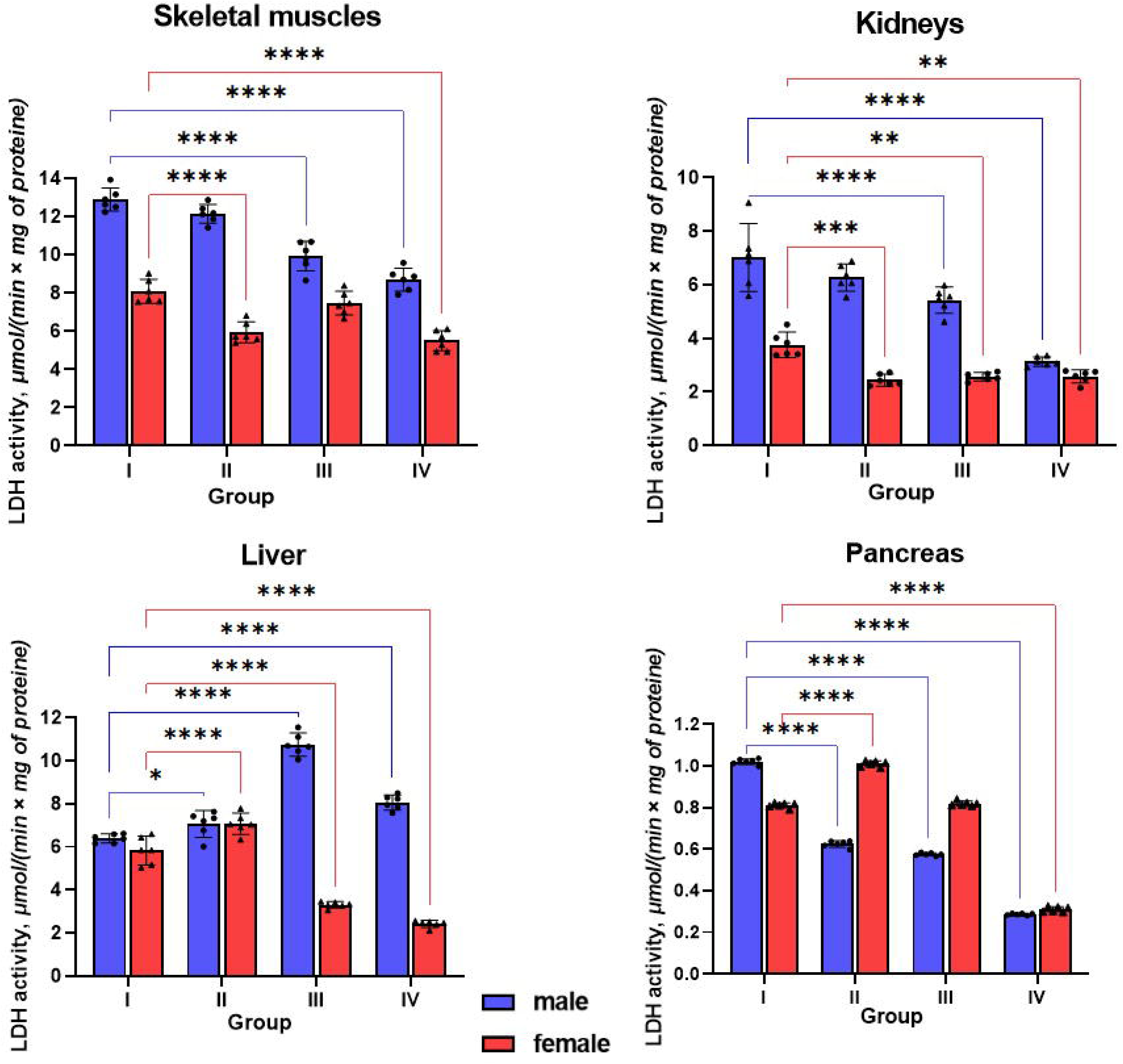
Effects of vanadium citrate on lactate dehydrogenase (LDH) activity in different tissues of male and female Wistar rats. Panels show specific LDH activity (μmol/(min × mg of protein)) in tissues. Animals were divided into four groups: I – control, II – 3 μg/kg body weight, III – 12.5 μg/kg body weight, IV – 50 μg/kg body weight vanadium citrate. Values are presented as mean ± 95% confidence interval (CI), n = 6. Asterisks indicate statistically significant differences compared to the corresponding control group: *p < 0.05, **p < 0.01, ***p < 0.001, and **** p < 0.0001.

The results of two-factor analysis of variance revealed a statistically significant effect of both sex (F(1,40)= 574.08, p<0.001) and vanadium citrate dose (F(3,40)=63.99, p<0.001) on LDH activity in rat muscles. In addition, a statistically significant interaction effect between sex and vanadium citrate dose was established (F(3,40)=23.21, p<0.05).

The obtained ANOVA results confirm the multidirectional effect of vanadium citrate on LDH activity in muscles depending on sex and dose. The significant effect of the factor “vanadium citrate dose” is reflected in the decrease in LDH activity in females, as well as in the consistent decrease in LDH activity in males in all studied groups compared to the control. The effect of sex indicates significant differences in the basal level of LDH activity in the muscles of females and males of the control group, which emphasizes these sex differences. The presence of a significant interaction effect between sex and dose of vanadium citrate indicates that the response of LDH activity to different doses of the drug is significantly different in females and males. This indicates sex-dependent differences in the regulation of LDH and regulatory mechanisms that depend on sex hormones and other physiological features of the organism.

The assessment of the effect size for LDH activity in muscles revealed that the sex of the animal makes the greatest contribution to the variability of the enzyme. This factor has a very strong influence, explaining 93.49% of the variance in LDH activity 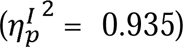, which is not explained by other factors in the model, indicating a significant role of gender in determining the variability of enzyme activity in muscle tissue. The effect of vanadium citrate dose was also significant, explaining 82.76% of the residual variance 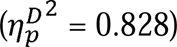. This indicates a significant effect of drug dose on LDH activity overall. In addition, the interaction effect between gender and vanadium citrate dose is moderate in magnitude, explaining 63.51% of the variance in LDH activity, indicating a significant contribution of gender-dependent dose responses to the variability of LDH activity 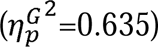.

For the remaining tissues, detailed results of the analysis of variance (two-way ANOVA) are presented in Table 1.

**Table 1.** Results of two-factor analysis of variance (ANOVA) of LDH activity in different tissues of rats under the influence of vanadium citrate (Partial eta-squared (η² ⍰) values represent the proportion of variance explained by each factor after accounting for other effects in the model and are therefore not additive).

| | Gender effect | Dose effect | Interaction effect | $\eta_p^{G^2}$<br>(%) | $\eta_p^{D^2}$<br>(%) | $\eta_p^{I^2}$<br>(%) |
| --- | --- | --- | --- | --- | --- | --- |
| **skeletal**<br>**muscles** | F(1,40)= 574.08<br>$p < 0.001$ | F(3,40)=63.99<br>$p < 0.001$ | F(3,40)=23.21<br>$p < 0.05$ | 93.49 | 82.76 | 63.51 |
| **kidneys** | F(1,40)= 290.85<br>$p < 0.001$ | F(3,40)= 46.16<br>$p < 0.001$ | F(3,40)= 21.74<br>$p < 0.001$ | 87.91 | 77.59 | 61.99 |
| **liver** | F(1,40)= 769.45<br>$p < 0.001$ | F(3,40)=49.87<br>$p < 0.001$ | F(3,40)= 226.69<br>$p < 0.001$ | 95.06 | 78.90 | 94.45 |
| **pancreas** | F(1,40)= 1115.75<br>$p < 0.001$ | F(3,40)= 6624.08<br>$p < 0.001$ | F(3,40)= 1529.57<br>$p < 0.001$ | 96.54 | 99.80 | 99.14 |

### Lactate dehydrogenase activity in the kidneys

The kidneys play a crucial role in maintaining glucose balance in the body, not only as an organ for gluconeogenesis, but also through the utilization of glucose as a metabolic substrate through glycolysis.

Determination of LDH activity in the kidneys is an important indicator of their condition. In the kidneys, under the influence of vanadium citrate, a decrease in LDH activity was observed in females of the experimental groups (Fig 3). Even the lowest dose led to a maximum decrease from 3.76 ± 0.36 μmol/(min×mg of protein) in the control (group I), to 2.44 ± 0.17 μmol/(min×mg of protein) at a vanadium citrate dose of 3 μgVCit/kg body weight in group II. Note that increasing the dose of vanadium citrate did not change the LDH activity in the experimental groups of females relative to each other, which allows us to conclude that the maximum decrease in LDH activity was achieved even at the lowest dose of vanadium citrate. The overall decrease in LDH activity in females in groups II, III and IV relative to the control was 35.1% (p < 0.001), 31.80% (p < 0.001) and 31.53% (p < 0.001), respectively. Males demonstrated a gradual decrease in LDH activity by 10.65% (p = 0.17) and 22.71% (p = 0.0087) and 55.39% (p < 0.001) relative to the control (group I - 7.01 ± 0.97 μmol/(min×mg of protein)) in groups II, III and IV, respectively (Fig 3). The decrease in LDH activity in the kidneys of female and male rats may indicate the inhibition of glycolysis under the action of vanadium citrate at all doses.

The obtained ANOVA results confirm the observed dose-dependent decrease in LDH activity in the kidneys under the action of vanadium citrate in both females and males (Table 1).

Similarly, the effect of different doses of vanadium citrate on the activity of LDH - a key enzyme of glycolysis and gluconeogenesis in the liver of rats was studied.

### Lactate dehydrogenase activity in the liver

The experimental data obtained by us demonstrate a significant difference in LDH activity in the liver of rats of both sexes depending on the dose of the studied compound. Small doses in females and males caused an increase in enzyme activity from 5.83 ± 0.51 (μmol/(min×mg of protein)) and 6.39 ± 0.16 (μmol/(min×mg of protein)) to 7.08 ± 0.38 (μmol/(min×mg of protein)) and 7.06 ± 0.47 (μmol/(min×mg of protein)) respectively (Fig 3). As we can see, the enzyme activity at a dose of vanadium citrate of 3 μgVCit/kg body weight does not differ between females and males, although in the control group males had higher LDH activity. At the same time the dose of 12.5 μgVCit/kg body weight led to a further increase in LDH activity in males by 68.30%, and a sharp decrease in activity in females by 43.26% compared to the control group. The highest dose of vanadium citrate (50 μgVCit/kg body weight) leads to a decrease in LDH activity in males (8.06 ± 0.27 (μmol/(min×mg of protein)) compared to animals that received the previous dose, but this value showed an increase of 26.22% compared to the control group. In females, the decrease in LDH activity continued, and at this dose it reached a level of 2.43 ± 0.13 (μmol/(min×mg of protein)), which is 58.38% less than the activity value in the control group.

These ANOVA results (Table 1) confirm that biological sex is the dominant factor determining the variability of LDH activity, while the interaction of sex with vanadium dose has an almost equally strong effect, which is critical for understanding the specific responses of the organism.

### Lactate dehydrogenase activity in the pancreas

In the pancreas, carbohydrate metabolism is regulated by the hormones insulin and glucagon, which are produced in the islets of Langerhans.

In the control group (Fig 3), LDH activity in the pancreas of females was 0.85±0.06 (μmol/(min×mg of protein)) and with vanadium intake at a dose of 3 μgVCit/kg body weight increased by 11.49% (vs I Group) to 0.95±0.12 (μmol/(min×mg of protein)) (p= 0.00216) (Fig. 3). At the same time, vanadium consumption at a dose of 12.5 μgVCit/kg body weight reduces LDH activity in the pancreas of females by 8.03% to 0.78±0.08 (μmol/(min×mg of protein)) without a significant difference (p= 0.5), a further increase in the dose to 50 μgVCit/kg body weight induces a further decrease in LDH activity by 63.91% (vs I Group) to 0.31±0.01 (μmol/(min×mg of protein)). All doses of vanadium citrate reduced LDH activity in the pancreas of males. If in the control this indicator was about 1.02±0.01 (μmol/(min×mg of protein)), then under the influence of the lowest dose of 3 μgVCit/kg body weight (Group II) it significantly decreased by 39.0% to 0.62 ±0.01 (μmol/(min×mg of protein)), and at a dose of 12.5 μgVCit/kg body weight (Group III) it decreased to 0.58 ± 0.01 (μmol/(min×mg of protein)) (≈ by 43.69% vs I Group).

The maximum three-fold decrease in LDH activity was observed in males receiving the highest dose of vanadium citrate (Group IV), where the indicator decreased to 0.29±0 (μmol/(min×mg of protein)) (p < 0.001 compared to Group I and other experimental groups).

This indicates the inhibition of glycolysis processes under the influence of all doses of vanadium citrate in the pancreas of males.

ANOVA indicators (Table 1) indicate fundamental and significant differences in the activity of the enzyme between females and males. Such high indicators were obtained due to the fact that the pancreas is the most sexually dimorphic organ in carbohydrate metabolism, and the response of LDH activity to vanadium citrate is almost completely modulated by both sex and the dose received.

Effect size analysis (partial ²) showed that sex-and dose-dependent responses were pronounced in all 4 tissues, but especially in the pancreas, indicating that biological sex is a major factor in the regulation of LDH in β-cells under the influence of vanadium citrate. The exceptionally high partial ² values for dose and interaction effects in the pancreas (99.80% and 99.14%, respectively) reflect the combination of a very large between-group difference (a three-fold decrease at the highest dose) with low within-group variability under controlled dosing and genetically homogeneous Wistar animals (n=6/group). Under these conditions, partial η² can approach unity even for group differences that are moderate in absolute terms; the corresponding F-statistics and 95% CIs shown in Fig 3 should therefore be interpreted alongside the effect-size estimates rather than in isolation.

### Analysis of the stability index

Analysis of the stability index (SI) revealed sex differences in the response to different doses (Table 2). In males, the lowest dose (3 μgVCit/kg body weight) provided the least deviation from control values, indicating the most stable metabolic profile at this dosage. In contrast, in females, the smallest deviations were observed at the intermediate dose (12.5 μgVCit/kg body weight), indicating their higher metabolic resistance to this level of stress. It is worth emphasizing that at the dose of 12.5 μgVCit/kg body weight, females demonstrate a significantly higher level of metabolic stability compared to males, in whom the SI at the same dose is 18.13% higher.

**Table 2.** Metabolic stability index (SI) in rats of both sexes depending on the dose of vanadium citrate.

|  | **Group II** | **Group II** | **Group IV** |
| --- | --- | --- | --- |
| **male** | 16.94±1.76 | 39.41±2.04 | 46.52±1.57 |
| **female** | 26.93±1.9 | 21.28±1.7 | 45.88±1.47 |

At the same time, it should be noted that at the lowest dose (3 μgVCit/kg body weight), the metabolic deviation in females remained higher than in males at their optimal dose (3 μgVCit/kg body weight), indicating a higher sensitivity of the metabolic system of females to the administration of vanadium citrate at the initial stages of the experiment.

In both sexes, the highest dose (50 μgVCit/kg body weight) resulted in the highest SI value (over 45 units), indicating pronounced metabolic disorders. This is consistent with the data on body weight dynamics (Fig. 2), where the most pronounced slowdown in animal growth rates was recorded at this dose. Thus, high values of the stability index reflect a significant restructuring of energy metabolism, which is visualized through the dynamics of body weight gain.

## Discussion

Many of the obtained patterns can be explained both by the action of vanadium and by the citrate form of vanadium. The vanadium citrate complex obtained by a two-step method (electric explosion) has high purity, bioavailability and stability compared to traditional vanadium salts [3].

It should be noted that citrate acts as an effective chelating (ligand) agent that encapsulates vanadium ions and changes their redox potential and transport pathways through cell membranes.

The changes in the activity of carbohydrate metabolism enzymes established in the study are based on the specific ability of vanadium and vanadium salts to mimic phosphate and modulate the functioning of key signaling cascades in the cell. The molecular effects of vanadium are predicted by the in-silico analysis of protein-metabolic interactions STITCH (Fig 4), which showed a possible high affinity of vanadate for tyrosine protein phosphatases (PTPs), in particular Ptpn1, Ptpn2, Ptprc, Ptprf, Ptprd and Ptprz1. By mimicking phosphate, vanadate blocks their active sites (score 0.88–0.96), which leads to a shift in the balance between phosphorylation and dephosphorylation of target proteins. Of particular importance is the inhibition of Ptpn1, a central negative regulator of insulin signaling. Since PTPs are closely related to Src family kinases (score up to 0.999), their blockade by vanadate produces a long-term increase in the level of phosphorylated proteins, with an increase in the insulin-like effect of vanadate.

**Fig 4.**
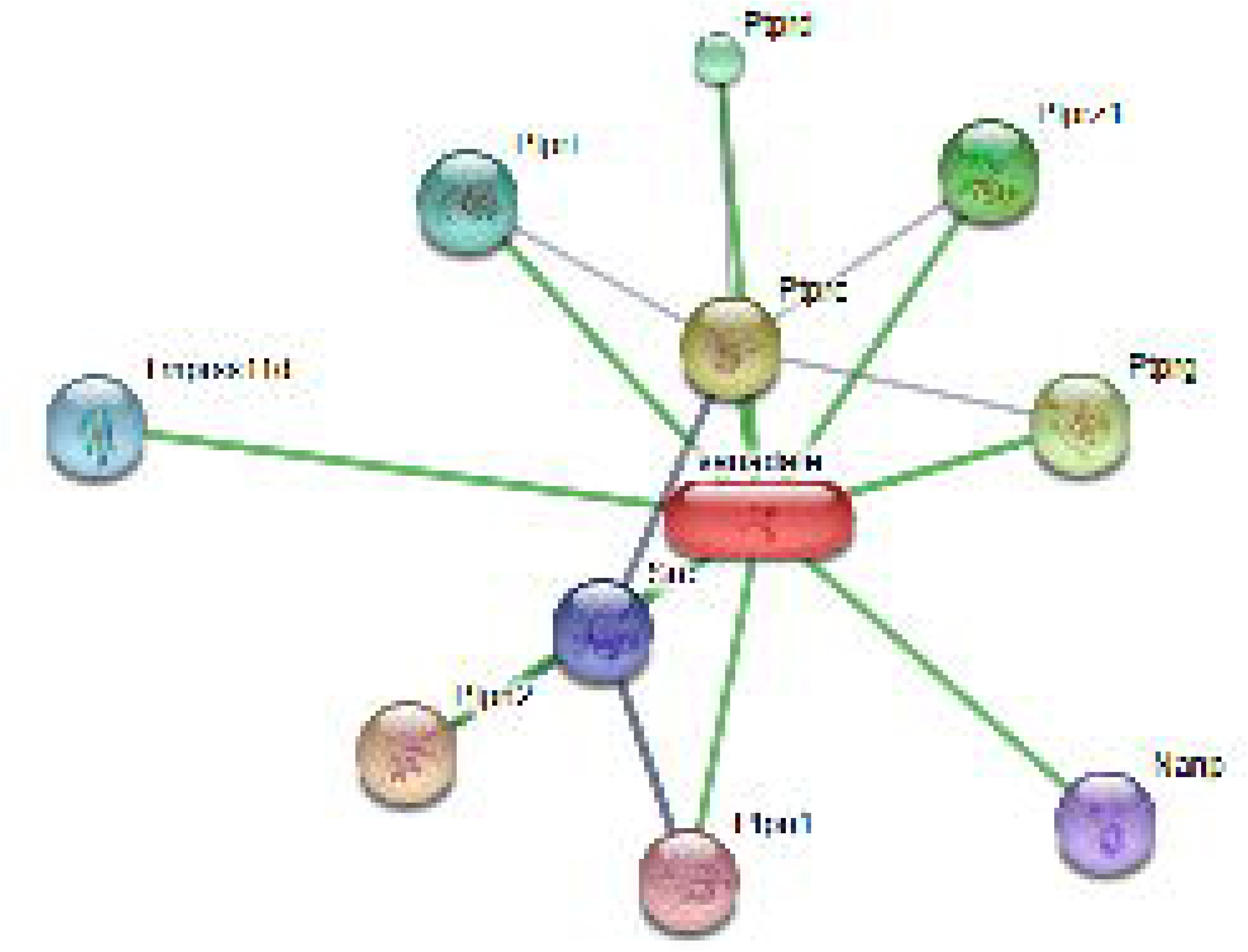
Protein-protein interactions associated with vanadate.

These mechanisms determine the nature of changes in LDH activity found in various rat tissues. LDH reflects the key balance between anaerobic glycolysis and mitochondrial oxidation and is sensitive to the cellular ratio *NADH/NAD^+^* [12].

At the same time, the citrate form likely allows for easier passage across lipid barriers (compared to inorganic salts), allowing for more efficient delivery of vanadium to its target molecules, particularly protein tyrosine phosphatases (PTPs).

The effect of citrate on LDH activity in cells is an example of a complex biochemical intervention involving both transport mechanisms and regulatory signaling pathways (Fig 5). Citrate, as a component of the complex, according to in silico predictions performed in STITCH, should ensure efficient delivery of vanadium into the cell, using the natural high-affinity sodium/citrate cotransporter Slc13a5, since the interaction coefficients of this protein with citrate (0.995) and sodium (0.985). Thanks to Slc13a5, citrate (and vanadium citrate in particular) may enter directly into the cytosol, where its further fate affects key carbon fluxes that have a direct connection with LDH. In the cytosol, elevated citrate concentrations may function as allosteric inhibitors of phosphofructokinase (PFK), a key regulator of glycolysis. If PFK-1 inhibition occurs under the present experimental conditions, it could reduce glycolytic flux and consequently decrease pyruvate availability for LDH. Thus, the citrate component of the complex may indirectly reduce LDH activity by reducing substrate availability.

**Fig 5.**
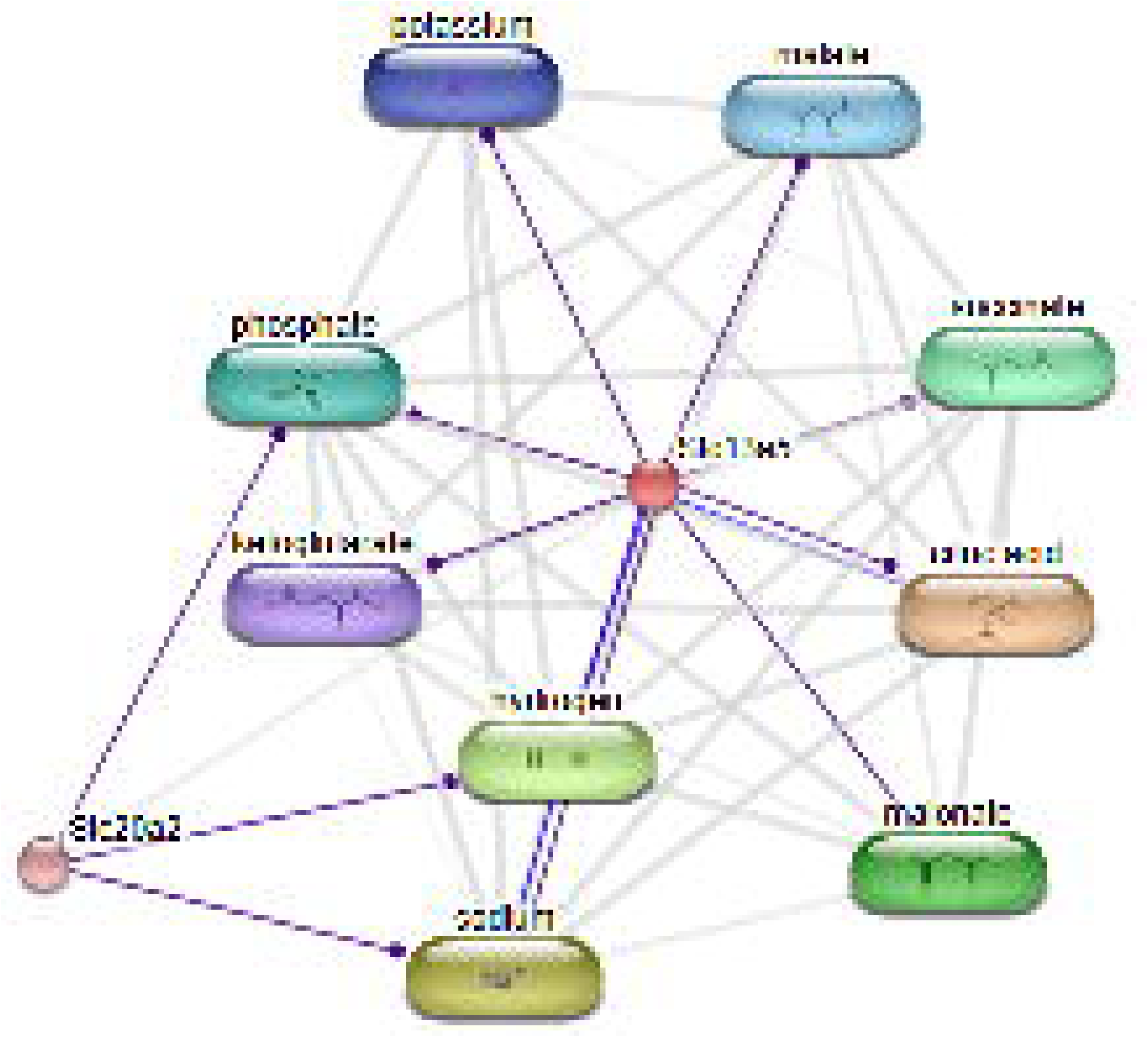
Molecular interactions of Slc13a5 as a potential pathway for the citrate ligand of vanadium to affect mitochondrial oxidation.

Since, in addition to the direct effect of citrate on glycolysis, the vanadium component exerts a stronger systemic and regulatory effect, and the insulin-like effect contributes to an overall increase in glucose metabolism, increasing the flow of pyruvate to the mitochondria for complete oxidation, rather than for reduction to lactate by LDH. In this context, the mediated interaction between Slc13a5 (citrate transport) and Slc20a2 (sodium-phosphate symporter) with a moderate index (0.665) also becomes important. This relationship, probably through shared sodium gradient, suggests a possible effect of vanadium on cellular phosphate homeostasis. Changes in phosphate concentration may directly affect the energy state of the cell and the rate of glycolytic reactions that regulate LDH activity.

Furthermore, the interaction network provided shows that citrate and other Krebs cycle metabolites (ketoglutarate, malate, succinate) interact strongly with hydrogen (0.927–0.951), highlighting the sensitivity of these pathways to cellular pH.

Since LDH is activated under acidotic conditions, any stabilization or change in pH induced by vanadium or citrate metabolism may indirectly affect its activity. Thus, the effect of vanadium citrate on LDH in different tissues (e.g., liver, where lactate oxidation dominates, or muscle, where it is generated) is dual: it is regulated both directly by citrate through substrate limitation and by vanadium through potent hormonal signaling and regulation of mitochondrial/energy balance.

This may explain the high efficacy of low doses of LDH that we observed (Fig 6).

**Fig 6.**
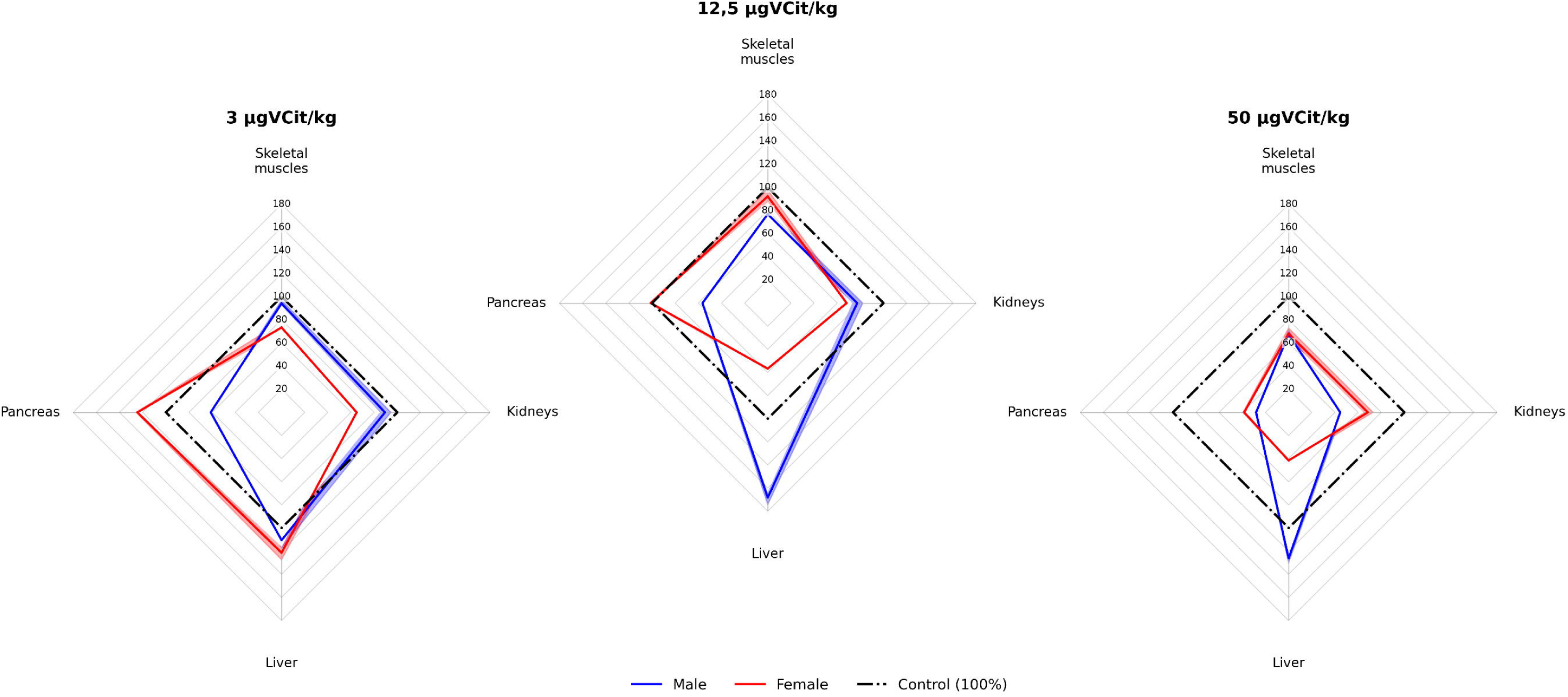
Radar plots of the stability index (SI) across different tissues in male and female Wistar rats under various vanadium citrate dosages.

These results are consistent with the radar plot analysis, which demonstrated more balanced tissue responses at these doses.

Although vanadium is known for its ability to accumulate with prolonged administration, the absence of changes in the female group (12.5 μg VCit/kg body weight) indicates the formation of metabolic adaptation. It is likely that at this dose the rate of accumulation of the complex in the tissues of females does not exceed the threshold of homeostatic correction, which indicates the safety of this concentration as a potential therapeutic dose. The use of young animals (6–11 weeks during the experiment) made it possible to record the moment of formation of different sensitivity to the dose in males and females, which would be more difficult to do in old animals with an already fading hormonal profile. Since animals of this age are in the phase of active weight gain, this made it possible to trace deviations in the growth rates of females and males when using different concentrations of vanadium citrate. Comparative analysis shows that sexual dimorphism determines a different metabolic response to the administration of vanadium citrate. In females, the differences in weight gain between the experimental groups were less contrasting, while in males, a significant gap was observed between the control group and the group receiving the highest dose (50 μg VCit/kg body weight).

A probable explanation for the observed slowdown in body weight gain, especially at high doses, is an active restructuring of metabolism towards energy optimization. Since the animals did not show signs of intoxication, it can be concluded that the used dosages of vanadium citrate are safe and are within the physiologically tolerable values for the rat organism.

Importantly, the stability index analysis does not support the existence of a single universal optimal dose. Instead, the data indicate a clear sex-dependent pattern, with males exhibiting the most stable response at lower doses and females at intermediate doses.

This emphasizes the importance of sex as a critical factor in the study of the metabolic effects of vanadium and indicates the presence of a narrow sex-specific therapeutic “window”. The stability index serves as an integrative marker; lower SI values in males at low doses and in females at intermediate doses reflect a state of “homeostatic optimization”. This indicates that at these specific doses, vanadium citrate modulates LDH activity, increasing aerobic efficiency without triggering systemic stress. The modest reduction in body weight gain at the highest dose (50 μg VCit/kg body weight) correlated with the largest deviation in the stability index and the most pronounced inhibition of LDH in all tissues. This reflects a metabolic shift in which cellular energy is directed toward intensive adaptation and regulation of the redox state rather than rapid biomass accumulation.

Overall, the observed changes in LDH activity reflect a multilevel effect of vanadium citrate, including inhibition of tyrosine phosphatases CPTPs) and enhancement of insulin-like signaling [13], as well as changes in redox state, a possible shift in the relative contribution of LDH-A/LDH-B isoforms (not directly assessed in this study), the influence of oxidative stress, and sex-dependent effects of estradiol and androgens (Fig 7).

**Fig 7.**
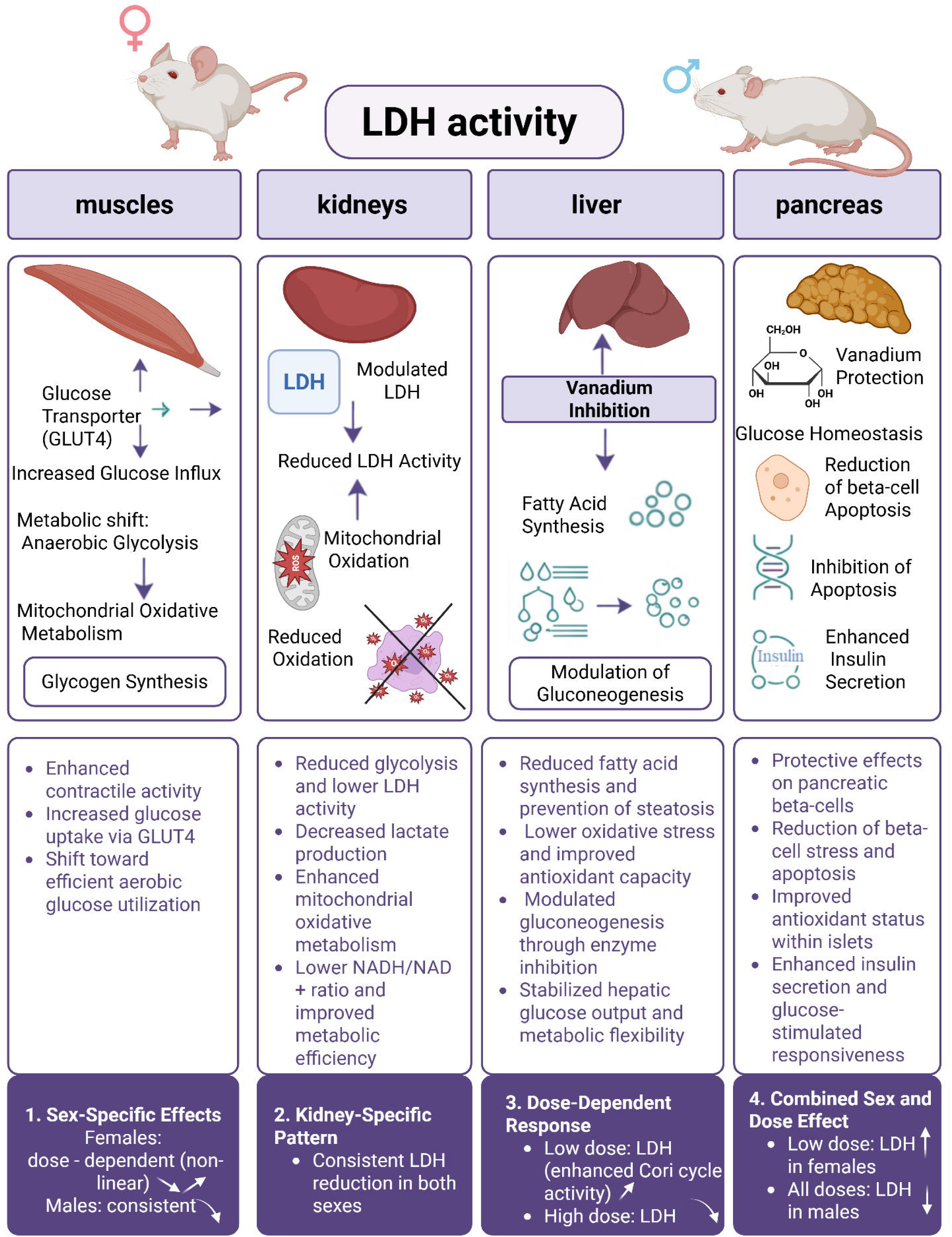
Schematic of tissue-and sex-specific metabolic effects of vanadium citrate in rats (Created with BioRender.com).

In skeletal muscle, where LDH-A is generally considered the predominant isoform, vanadium citrate induced opposite sex responses: females showed a dose-dependent decrease and recovery of LDH activity (prior dose 12.5 μg VCit/kg body weight), consistent with estrogen-mediated stimulation of the glycolysis pathway [14].

In contrast, males showed a sustained decrease in activity, reflecting a significant shift in metabolism toward the oxidative pathway and a weaker response to the insulinomimetic effects of vanadium [15]. The decrease in LDH activity in skeletal muscle may indicate that pyruvate is more efficiently directed to the mitochondria for aerobic oxidation (via the Krebs cycle) rather than being converted to lactate (via LDH).

The observed decrease in LDH activity may indicate a metabolic switch to oxidative pathways, which potentially increases the efficiency of glucose utilization under the influence of vanadium citrate. Thus, the decrease in LDH activity is a possible physiological consequence of the fact that muscles under the influence of vanadium citrate use glucose (and probably oxygen) more efficiently for energy, and therefore have less need for the anaerobic pathway (which requires high LDH activity). In particular, in [16] it was reported that it promotes muscle contractile activity, which in turn leads to increased glucose uptake, as a result of which the need for insulin disappears. Renal LDH in animals of both sexes decreased at all doses, which may be due to high mitochondrial activity of renal tissue. This decline is consistent with, though not direct evidence for, inhibition of the pyruvatogenic LDH-B isoform, which is known to be highly sensitive to oxidative modification, as well as a decrease in cytosolic NADH and, as a result, a violation of the inclusion of pyruvate in the tricarboxylic acid cycle [17]. This is consistent with the known high sensitivity of enzymes (including LDH-B) to oxidative modification, which may be enhanced by higher doses of vanadium [2].

This decrease in renal LDH activity may reflect an adaptation of glucose-pyruvate fluxes, favoring aerobic oxidation in tissues with high mitochondrial density.

Hepatic LDH showed a nonlinear, U-shaped response curve. Low doses caused a modest increase in activity, which is a sign of increased glycolysis and activation of the Cori cycle (compensatory gluconeogenesis). While higher doses might exert an inhibitory effect on the enzyme, possibly due to a dose-dependent modulation of the cellular redox state [19]. This pattern is broadly consistent with reports on other vanadium complexes: hepatic protection against lipid peroxidation and endoplasmic reticulum stress has been described for vanadium(IV)-chlorodipicolinate at moderate doses [20, 21], while pancreatic β-cell responses to vanadium have similarly been shown to shift from a protective, glucose-toxicity-relieving effect at controlled exposure toward dysregulation at higher or prolonged doses [24]. The tissue-and dose-dependent trajectory observed here for LDH activity in liver and pancreas therefore parallels the broader biphasic pattern reported for vanadium compounds across these organs, rather than representing an isolated finding specific to the citrate ligand. The stimulatory effect of low concentrations of vanadium citrate on LDH activity in both sexes and the inhibitory effect of high concentrations reflect the general trend, and the quantitative differences in the enzyme response depending on the sex are also statistically significant, reflecting the high metabolic flexibility of the liver in maintaining glucose homeostasis. Hormonal differences can affect fat metabolism and gluconeogenesis in the liver and kidney, which directly modulates the need for LDH.

The pancreas is characterized by low basal LDH activity, since β-cells predominantly oxidize pyruvate in the mitochondria to ensure insulin secretion [22]. In our study, low doses in females increased LDH activity in the pancreas, enhancing lactate generation, which may be potentially reduces the secretory capacity of β-cells by diverting pyruvate from the mitochondria [22]. A dose of 12.5 μg VCit/kg body weight did not result in changes in LDH activity in the pancreas, which may be related to reaching a homeostatic equilibrium level, in which pyruvate metabolic fluxes maintain their physiological orientation toward aerobic oxidation.

However, high doses in females and all doses in males resulted in inhibition of the enzyme, reflecting significant changes in the enzymatic regulation of pyruvate metabolism in β-cells, which are highly sensitive to vanadium exposure [23]. Increased LDH in β-cells is functionally unfavorable for function (removal of pyruvate from mitochondria) [19, 22], and a corresponding decrease may be protective (combating glucose toxicity [23]), but only up to a certain point (where the toxic effect of vanadium begins). Accordingly, the extremely high size of the sex effect for pancreatic LDH activity (Table 1) reflects endocrine-specific metabolic dimorphism of β-cells, rather than a generalized systemic effect. Therefore, under high-dose conditions (Group IV), the decrease in LDH activity in the liver and pancreas may be due not only to the inhibitory effects of supra-optimal concentrations of vanadium, but also to the combined effects of vanadium (which inhibits gluconeogenesis/glycolysis) and the high citrate content, which further inhibits PFK-1, slowing glycolysis and, consequently, reducing the need for LDH. Since citrate (especially at high concentrations) is an allosteric inhibitor of the key enzyme of glycolysis, phosphofructokinase-1 (PFK-1), this is a unique effect of the citrate form.

Taken together, the observed changes in LDH activity are consistent with a hypothesized dual mechanism combining insulin-mimetic action of vanadium via inhibition of protein tyrosine phosphatases and citrate-mediated allosteric inhibition of PFK-1. This interpretation is supported by *in silico* interaction data (STITCH) and the observed pattern of LDH activity, but has not been directly verified; confirmation would require measurement of PFK-1 activity, tissue pyruvate/lactate pools, and mitochondrial oxygen consumption rate.

One limitation of this study is the lack of control for the stage of the estrous cycle in females. Thus, variability in the stages of the estrous cycle could have partially influenced the obtained values of LDH activity and contributed to the intragroup variability in females. This should be taken into account when interpreting sex differences and in future studies, control or stratification by cycle stage should be provided. Isoform-specific LDH activity (LDH-A vs LDH-B) was assessed indirectly through total spectrophotometric activity; native PAGE or western blotting would be required to directly confirm isoform-level shifts.

In addition, the proposed dual mechanism of action — PFK-1 inhibition by the citrate ligand and redirection of pyruvate toward mitochondrial oxidation — is inferred from LDH activity patterns and *in silico* protein-interaction predictions rather than direct biochemical measurement. PFK-1 activity, tissue pyruvate and lactate concentrations, and mitochondrial oxygen consumption were not assessed in this study and represent a priority for follow-up work.

## Conclusions

The detected changes in LDH activity in key metabolic tissues of rats indicate multilevel, sex-, tissue-and dose-dependent effects of the vanadium citrate complex. The obtained results are consistent with a hypothesized synergistic dual mechanism, which may explain its high bioavailability and pronounced metabolic effect, pending direct biochemical confirmation (see Limitations). On the one hand, the vanadium component of the complex implements a systemic regulatory effect through the imitation of the phosphate group and inhibition of protein-tyrosine phosphatases, in particular PTPn1, which leads to a long-term increase in the level of phosphorylation of signaling proteins and strengthening of the insulin cascade. On the other hand, the citrate ligand may contribute to metabolic regulation through its known ability to inhibit PFK-1 allosterically; however, this mechanism was not directly assessed in the present study.

At all doses, vanadium citrate led to a slight decrease in the rate of weight gain. The lowest intensity of gain was observed in animals receiving the highest concentrations of vanadium. It decreased by 30.1% in the female group and by 35.3% in the male group. However, despite this decrease, the general condition of the animals remained without signs of physiological disorders or pathological deviations in behavior.

Two-way analysis of variance confirmed a statistically significant effect of both dose and sex of animals, as well as their interaction, on LDH activity in all tissues studied, including skeletal muscle, kidney, liver and pancreas. In males, the effect of vanadium citrate was manifested mainly in a decrease in LDH activity (except in the liver), which may reflect a reorientation of energy metabolism from anaerobic glycolysis to oxidative pathways with a more efficient direction of pyruvate into mitochondrial oxidative metabolism. In contrast, in females, the effect of the complex was clearly dose-dependent: at low doses, an increase in the activity of carbohydrate metabolism enzymes in the liver and pancreas was observed, which may indicate the activation of compensatory glycolysis and the Cori cycle, while at high doses, LDH inhibition was observed, possibly related to allosteric PFK-1 inhibition by the citrate ligand (not directly measured in this study) and manifestations of vanadium toxicity. The detected sex differences are consistent with the mediated role of sex hormones, in particular estrogens, which are known to increase the efficiency of aerobic oxidation and metabolic plasticity of tissues. The tissue-specific nature of the changes in LDH activity reflects the balance between the adaptive and potentially harmful effects of the vanadium citrate complex. In skeletal muscle, both in men and women, the reduction in LDH activity can be considered as a sign of a more energy-efficient and metabolically “healthy” state, in which glucose is predominantly used by aerobic pathways in the presence of an adequate oxygen supply. In highly oxidative organs, such as the kidney and liver, the action of vanadium citrate is formed as a compromise between insulinomimetic stimulation of metabolic pathways at low doses and direct inhibition of glycolysis. The response of the pancreas is particularly indicative with a narrow therapeutic window: the increase in LDH activity observed at low doses in females is potentially undesirable because it diverts pyruvate from mitochondrial oxidation in β-cells, whereas the inhibition of LDH observed at high doses or in men at all levels of exposure may have a protective effect, reducing glucose toxicity before systemic vanadium toxicity occurs. Thus, the observed synergistic pattern of the vanadium and citrate components of the complex at the level of regulation of protein tyrosine phosphatases and key enzymes of glycolysis provides a plausible molecular framework for its metabolic effects, which warrants direct biochemical validation — in particular, measurement of PFK-1 activity, tissue pyruvate/lactate levels, mitochondrial oxygen consumption, and isoform-specific LDH-A/LDH-B activity via native PAGE or western blotting. The results obtained emphasize the potential of vanadium complexes as metabolic modulators with tissue-, dose-, and sex-specific effects, which is relevant for establishing safe physiological dosing thresholds and understanding early, subclinical metabolic responses to vanadium exposure in a healthy organism, particularly in the liver and pancreas. SI show that for males the most stable (physiological) dose is 3 μg VCit/kg body weight (SI = 16.94), while for females it is 12.5 μg VCit/kg body weight (SI = 21.28). This highlights the need for a personalized approach to vanadium dosing depending on gender.

Future work should directly address the mechanistic limitations noted above by combining LDH activity measurements with PFK-1 assays, tissue pyruvate/lactate quantification, mitochondrial oxygen consumption analysis, and isoform-specific LDH separation.

This work represents the first part of a comprehensive study of the metabolic effects of vanadium. In the next publication, glucose-6-phosphate dehydrogenase activity and redox mechanisms that determine antioxidant and adaptive tissue responses will be analyzed in detail.

## Supporting information

**S1 Data. Supporting raw data.** All primary data underlying the findings are available at the following Zenodo repository: https://doi.org/10.5281/zenodo.19420548.

